# Molecular Basis for the Adaptive Evolution of Environment Sensing by H-NS Proteins

**DOI:** 10.1101/2020.04.21.053520

**Authors:** Xiaochuan Zhao, Vladlena Kharchenko, Umar F. Shahul Hameed, Chenyi Liao, Franceline Huser, Jacob M. Remington, Anand K. Radhakrishnan, Mariusz Jaremko, Łukasz Jaremko, Stefan T. Arold, Jianing Li

**Author notes:** Correspondence to: JL, STA or LJ.

## Abstract

The DNA-binding protein H-NS is a pleiotropic gene regulator in gram-negative bacteria. Through its capacity to sense temperature and other environmental factors, H-NS allows pathogens like Salmonella to adapt their gene expression, and hence toxicity and biological responses, to their presence inside or outside warm-blooded hosts. To investigate how this sensing mechanism may have evolved to fit different bacterial lifestyles, we compared H-NS orthologs from bacteria that infect humans, plants, and insects, and from bacteria that live on a deep-sea hypothermal vent. The combination of biophysical characterization, high-resolution proton-less NMR spectroscopy and molecular simulations revealed, at an atomistic level, how the same general mechanism was adapted to specific habitats and lifestyles. In particular, we demonstrate how environment-sensing characteristics arise from specifically positioned intra- or intermolecular electrostatic interactions. Our integrative approach clarified the mechanism for H-NS–mediated environmental sensing and suggests that it resulted from the exaptation of an ancestral protein feature.

## INTRODUCTION

The histone-like nucleoid-structuring (H-NS) protein is a central controller of the gene regulatory networks in enterobacteria (*1*). H-NS inhibits gene transcription by coating and/or condensing DNA; an environment-sensing mechanism allows H-NS to liberate these DNA regions for gene expression in response to physicochemical changes (*2–4*). H-NS preferentially binds to AT-rich sequences, which enables its dual role in (i) the organization of the bacterial chromosome and (ii) the silencing of horizontally acquired foreign DNAs (*5–8*). The latter mechanism allows bacteria to assimilate foreign DNAs, which, however, are only expressed as a last resort in case of acute threats or stresses (*8*). Thus, H-NS plays a crucial role in the adaptation, survivability, and antibiotic resistance of bacteria. Given the growing threat of multidrug resistance, H-NS has attracted increasing research interest, with a particular focus on elucidating the molecular mechanisms of adaptive evolution (*9–11*).

H NS possesses two dimerization domains (site1, residues 1-44; site2, resides 52-82; the numbering of *Salmonella typhimurium* is adopted throughout the text), and a C-terminal DNA-binding domain (DNAbd, residues 93-137) that is connected through a flexible region (linker, residues 83-92) to site2 (**Fig. 1**) (*5, 13–16*). The combination of site1 and site2 dimers allows HNS to form multimers for a stable concerted DNA association that results in gene silencing (*15*). In a previous study, we showed that site2 of *S. typhimurium* H-NS is the primary response element to temperature changes (*17*). Site2 unfolds at human body temperature, allowing the linker-DNAbd region to associate with site1 to adapt an autoinhibited conformation incapable of binding to DNA. Salinity and pH can also influence the stability of site2 dimers, and hence may also affect gene repression by H-NS (*17, 18*). Thus, the sensitivity of H-NS to temperature and other physiochemical changes allows human pathogens such as *S. typhimurium, Vibrio cholerae*, and enterohaemorrhagic *Escherichia coli* to sense when they enter a homothermic host and adapt their gene expression profiles accordingly.

**Figure 1.**
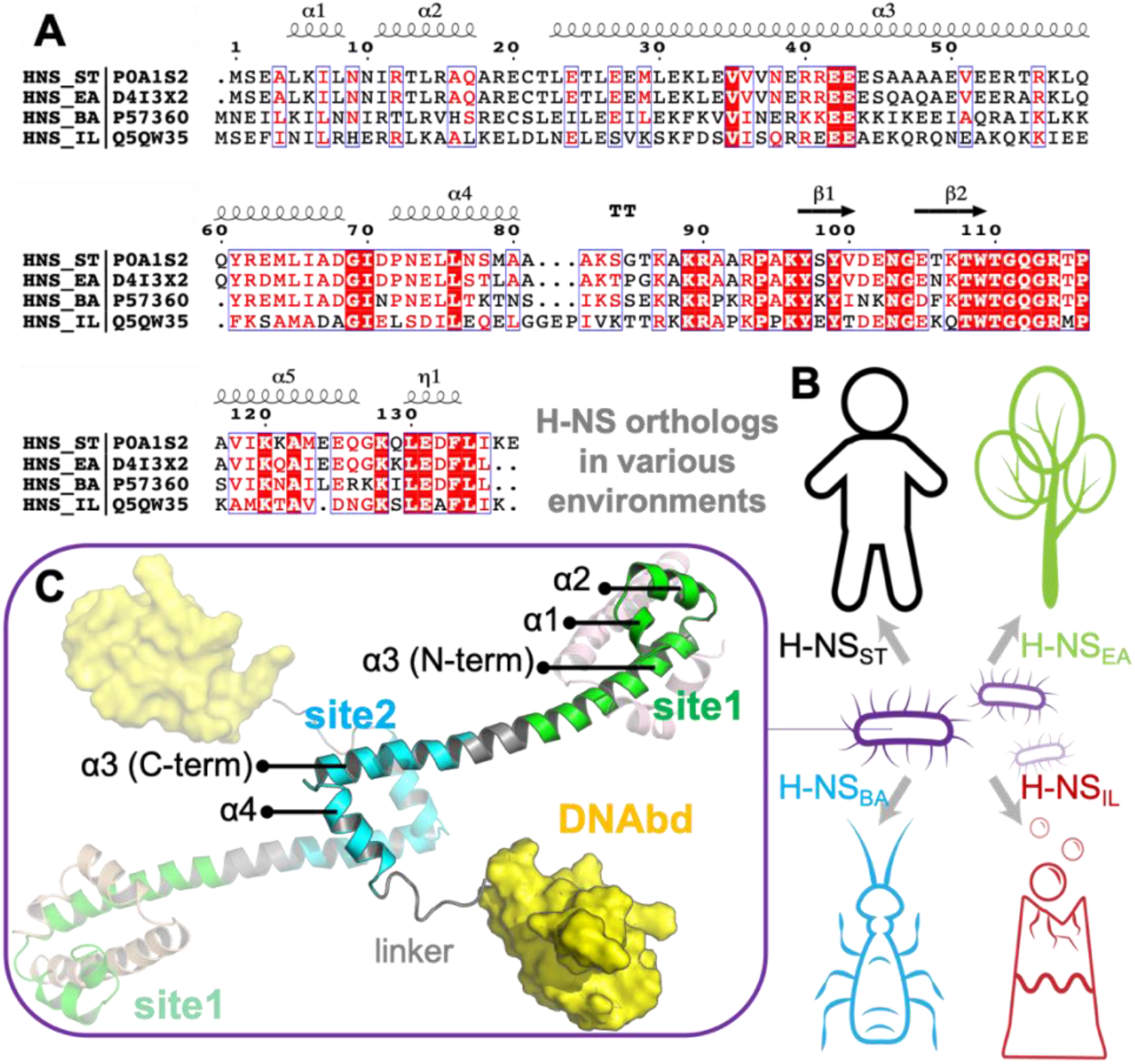
The sequence, structure, and habitat of selected H-NS orthologs. **(A)** Sequence alignment of HNS orthologs was performed with ESPript 3.0 (*12*): H-NS_ST_ (UniprotID: P0A1S2), H-NS_EA_ (UniprotID: D4I3X2), H-NS_BA_ (UniprotID: P57360), and H-NS_IL_ (UniprotID: Q5QW35). **(B)** Illustration of the environment of the selected orthologs. **(C)** The tetrameric H-NS_ST_ model we used for MD is shown with each domain labelled: site1 (green helices), site2 (cyan helices), linker (grey loop), and DNA-binding domain (yellow surface). The truncated site1 of flanking site1-dimerized chains are shown in magenta and pale orange. The tetramer H-NS_ST_ model was shown in cartoon, with each domain labelled: site1 (residues, 1-44), site2 (52-82), linker (83-92), and DNA-binding domain (93-137).

To date, studies to elucidate environment sensing of H-NS were almost exclusively conducted with proteins from two model systems, *S. typhimurium* (e.g. (*9, 19–21*)) and *E. coli* (e.g. (*18, 22–26*)), both of which infect humans. Yet, H-NS orthologs are also present in enterobacteria that do not have warm-blooded hosts, raising the question of what biological role H-NS plays in these species. Answering this question requires to determine the structural basis for environment-sensing in H-NS orthologs with drastically different lifestyles. However, the molecular dynamics and multidomain composition of H-NS hamper conventional structural analysis. Therefore, we combined large-scale molecular simulations and spectroscopic approaches to elucidate how environment-sensing by H-NS may have adapted in different species. This pluridisciplinary approach yielded an atomic-level understanding of how H-NS orthologs evolved specific residue substitutions to adapt environment-sensing to their bacterial habitats, and may open new avenues for strategies to combat antibiotic resistance.

## RESULTS

To investigate the adaptation of environment-sensing by H-NS, we selected four H-NS orthologs from ~3000 H-NS-like sequences available in the Uniprot database: 1) H-NS_ST_ from *S. typhimurium* (a pathogen of mammals), 2) H-NS_EA_ from *Erwinia amylovora* (a plant pathogen that infects apples and pears), 3) H-NS_BA_ from *Buchnera aphidicola* (an endosymbiont of aphids), and 4) H-NS_IL_ from *Idiomarina loiheinsis* (a free-living bacterium from a deep-sea hypothermal vent). H-NS_EA_ and H-NS_BA_ share more than 60% sequence identity with H-NS_ST_, whereas H-NS_IL_ is only 40% identical to H-NS_ST_ (**Fig. 1A**). Across the orthologs, the least conserved regions are residues 45–56 in α3, and residues 77–86 (end of α4 and beginning of the linker). The variable residues on α3 and linker may act as simple spacers (**Fig. 1**), suggesting α4 as a prime candidate for mediating adaptations to the environment.

### The site1 dimer is markedly more stable than the site2 dimer in the H-NS orthologs

H-NS_ST_ site1 and site2 form homodimers to enable H-NS multimerization in a head-to-head/tail-to-tail fashion (**Fig. 1C**) (*15*). In concert with the DNA interaction of the individual domains, this homooligomerization is required for tight DNA binding and hence gene repression. In our previous study, we showed that only H-NS_ST_ site2 dimers unfold and dissociated within a biologically relevant temperature range, whereas site1 dimers remain unaffected (*17*). The higher stability of the site1 dimer of H-NS_ST_ is explained by a substantially larger contact surface between the two monomers (ca. 3,300 Å_2_ compared to ca. 850 Å_2_ for site1 and site2, respectively, according to PDBePISA (*27*)).

To investigate whether this mechanism is conserved in other H-NS orthologs, we built homology models for H-NS_EA_, H-NS_BA_, and H-NS_IL_ using the crystal structure of the H-NS_ST_ site1-site2 fragment as a template (PDB ID: 3NR7) (*15*). Next, we constructed a tetrameric model as a minimal representation that conserves all features of the H-NS multimer. This tetramer contained two full-length H-NS monomers (residues 1-137, with templates PDB IDs: 3NR7 and 2L93) and two partial monomers, truncated before site2 (residues 1-52) (**Fig. 1C**). To probe differences in environmental responses of the orthologs, we first used conventional full-atom molecular dynamics (MD). We simulated all four tetramers (a ~100,000 atom system; see **Methods**) for 200 ns at three different conditions (0.15 M NaCl, 293 K; 0.50 M NaCl, 293 K / 20 °C; or 0.15 M NaCl, 313 K / 40 °C) (**Supplementary File 1A**).

The tetramer simulations at 0.15 M NaCl and 293 K produced a lower residue fluctuation level in site1 (local root-mean-squared fluctuation, RMSF 0.4 to 1.9 Å) than in site2 (local RMSF 0.5 to 4.4 Å) for all four orthologs (**Supplementary File 1B**). The higher stability of the site1 dimer is explained by the generally higher number of nonpolar contacts than in the site2 dimer (**Fig. 2, Figure Supplement S1A**). These contacts involved conserved hydrophobic amino acid residues, notably L5 (or I5) and L8 of α1, L14 of α2, and L23, L26, V36, and V37 (or I37) of α3 (**Fig. 2**). These interactions remained formed in all site1 dimers in our tetramer simulations (at 0.15 M NaCl at 293 K) and tetramer simulations at higher salinity (at 0.50 M NaCl at 293 K) or higher temperature (at 0.15 M NaCl at 313 K). Hence, we found that the stability of the site1 dimers resulted mainly from strong and conserved nonpolar packing. Collectively, these data conclude that the mechanism observed for H-NS_ST_ — where site1 remains stable, and the site2 stability is affected by the environment — is conserved in H-NS_EA_, H-NS_BA_, and H-NS_IL_.

**Figure 2.**
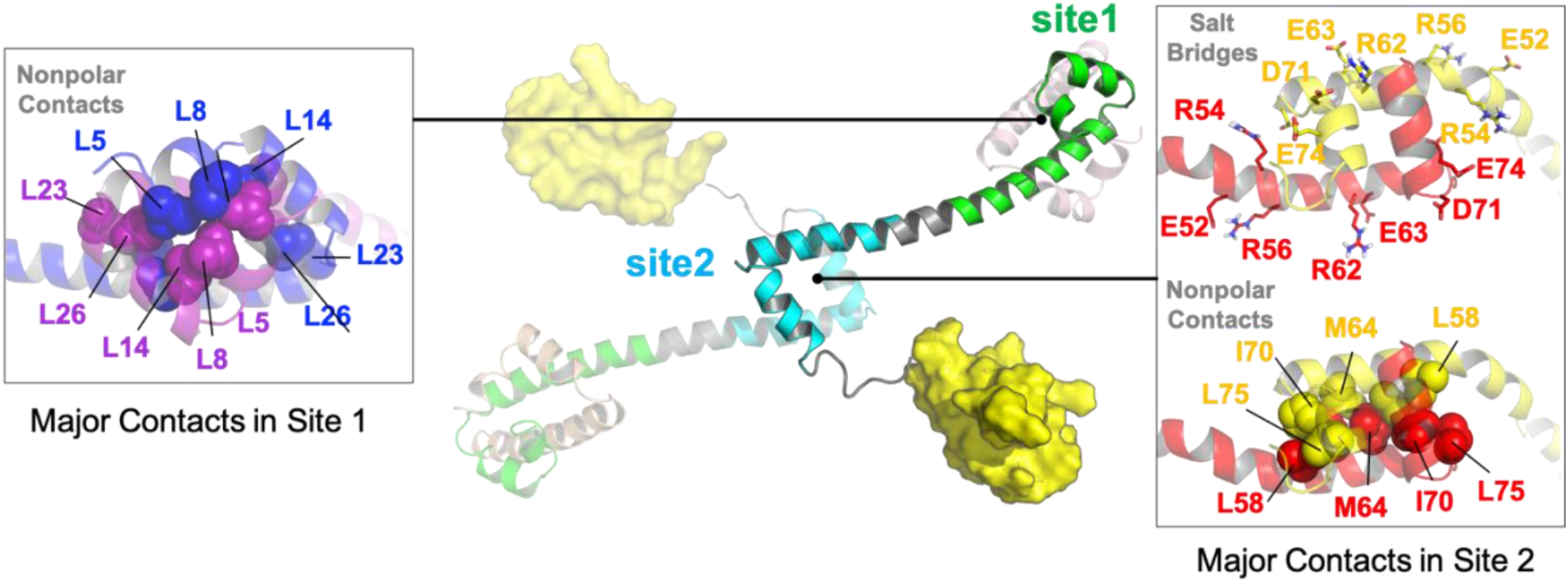
Contacts in site1 and site2 dimers in H-NS_ST_. Hydrophobic contact residues are shown as sphere models, and polar contact residues as stick models. Two protein chains forming the dimer are color-coded. For additional details see **Figure Supplement S1.**

### Variations in the site2 sequence alter the sensing sensitivity of H-NS orthologs

Compared to site1 dimers, site2 dimers harbor fewer nonpolar contacts, only involving residues L58 (or I58), Y61 (or F61), M64 (or A64), I70, and L75 (or I75) (**Fig. 2; Figure Supplement S1A**). Hence, while site1 dimerization was largely maintained by nonpolar packing, site2 dimerization was strongly driven by electrostatic interactions from selective salt bridges. MD simulations revealed that these salt bridges were in a dynamic equilibrium between forming, breaking, and rearranging. These salt bridges were either formed *in cis*, within the site2 monomer (e.g., E52-R56 and R62-E63 in H-NS_ST_) or *in trans*, between two monomers in the site2 dimer (e.g., R54-D71’, R54-E74’ and K57-D68’ in H-NS_ST_; where the apostrophe denotes residues from the second chain; illustrated in **Fig. 3**). In addition to substitutions that delete (E52A in H-NS_BA_; E63S and D68A in H-NS_IL_) or weaken (E63Q in H-NS_EA_; D71N in H-NS_BA_) these salt bridges, our simulations showed different levels of site2 salt bridge stability among orthologs (**Fig. 3, Supplementary File 1C**): (i) The intermonomer salt bridge R/K54-E/D74’ was stable in all our simulations at 293 K and 0.15 M NaCl, but less likely to form at an increased temperature (313 K / 40 °C) or salinity (0.50 M NaCl), suggesting that this salt bridge is involved in environmental sensing (**Fig. 3A**). (ii) Absent in H-NS_IL_, the inter-monomer salt bridge K57-D68’ remained formed during all our simulations of H-NS_ST_, H-NS_EA_, and H-NS_BA_, indicating a ‘housekeeping’ role for the stability of the site2 dimer in all orthologs except for H-NS_IL_ (**Fig. 3B**).

**Figure 3.**
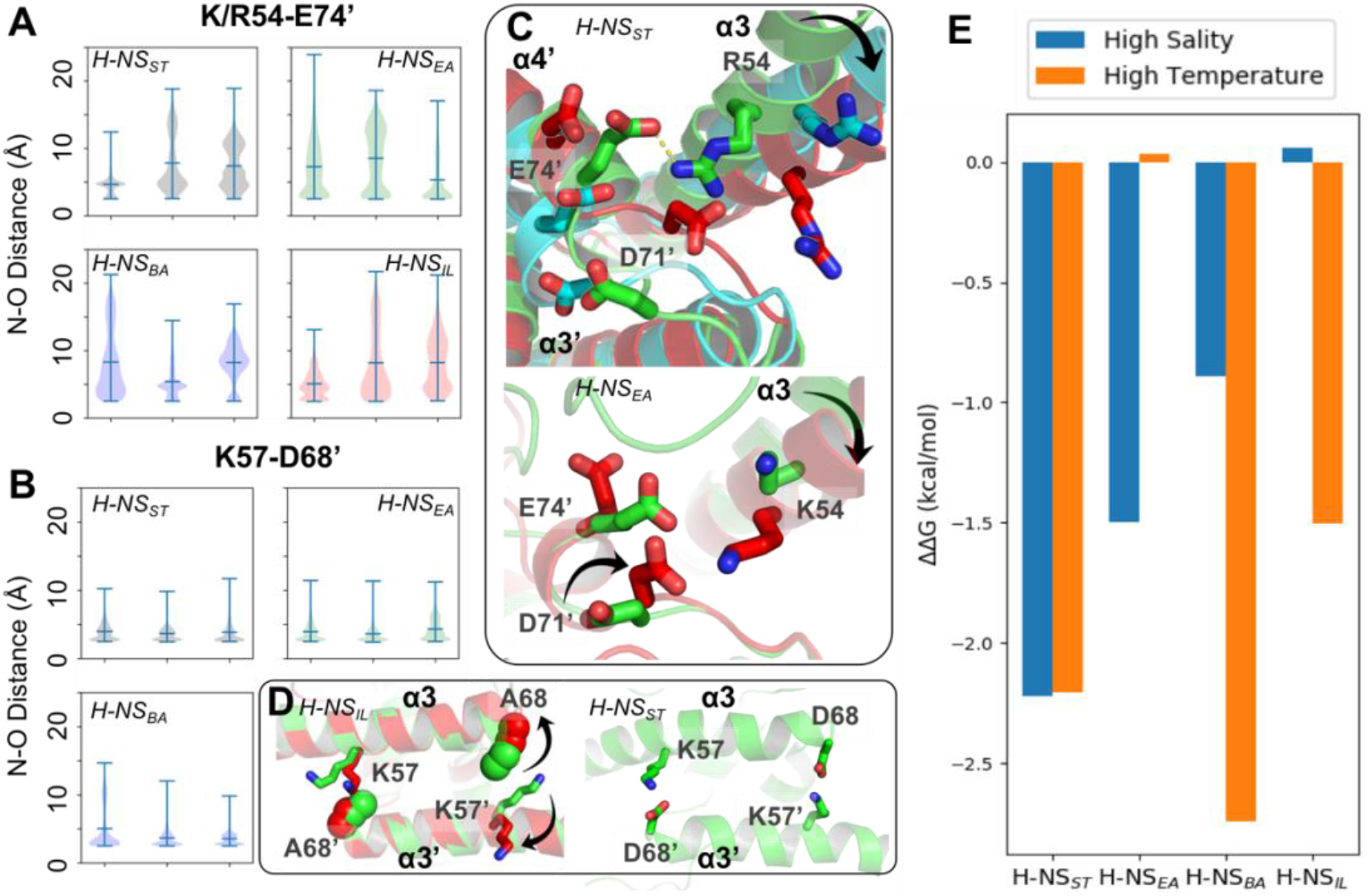
Conserved inter-dimer salt bridges observed in MD simulations. **(A-B)** Violin plots of the distance between the nearest sidechain nitrogen atom of lysine/arginine and the sidechain oxygen atom of aspartic/glutamic acid in the salt bridge. Each subplot shows the results obtained at 293 K, 0.15 M NaCl (left), 313 K, 0.15 M NaCl (middle), and 293 K, 0.50 M NaCl (right). Each “violin” displays the mean value (the bar in the center of the violin), the range (the stretched line), and the distribution of the distance (kernel density on the side). As we use the numbering of H-NS_ST_, there are position shifts in H-NS_BA_ and H-NS_IL_: R54 to R53, K57 to K56 and D68 to D67 in H-NS_BA_; R54 to K53 in H-NS_IL_. **(C)** Final snapshots of the R54-E74’ salt bridge in H-NS_ST_ and K54-D71’ in H-NS_EA_ (s. **Figure Supplement S2** for additional details). Color scheme of the cartoon: 293 K, 0.15 M NaCl (green), 313 K, 0.15 M NaCl (red), and 293 K, 0.50 M NaCl (cyan). **(D)** Final snapshots of the K57-D/A68 contact in H-NSST and H-NSIL. Same color scheme as (C). **(E)** Free energy changes as a result of increased salinity or temperature, according to the PMFs calculated from umbrella sampling. (s. **Figure Supplement S3** for additional details).

Our simulations show how specific protein dynamics might modulate the ortholog’s response to salinity or temperature. For example, we observed increased bending of the α3 backbone (annotated by the black arrow in **Fig. 3C**) at high temperature (313 K) or high salinity (0.50 M NaCl) (**Figure Supplement S2**). Although α3 bending occurred in all orthologs, it only significantly affected the site2 dimer of H-NS_ST_ by separating R54 from E74’ or D71’, suggesting that this mechanism contributed to the salt and temperature sensitivity of H-NS_ST_ site2, whereas it was not strong enough to significantly affect site2 stability in other orthologs.

Another example was given by H-NS_EA_, where an alternative R54-D71’ salt bridge formed whenever the R54-E74’ contact was broken at 313 K. This alternative R54-D71’ salt bridge stabilized the H-NS_EA_ site2 dimer at the higher temperature, suggesting that this compensatory mechanism resulted in a decreased sensitivity to temperature (**Fig. 3C**). H-NS_IL_ provided a final example for a specific response. Compared with the R54-E74’ salt bridge (**Fig. 3A**), the K57-D68’ salt bridge only varied slightly in all our simulations (**Fig. 3B**). However, the substitution D68A in H-NS_IL_ supplanted the electrostatic interaction with a nonpolar interaction, which was broken at 313 K in our simulations (**Fig. 3D**). This effect suggested that H-NS_IL_ had a reduced sensitivity to salinity, while remaining sensitive to temperature.

To complement the dynamics of H-NS orthologs from our conventional MD simulations, we used extensive simulations with umbrella sampling to quantitate the overall site2 stability. We calculated the potential of mean force (PMF) for site2 dimer dissociation (residues 50-82, ca. 46,000 atoms) of the four H-NS orthologs for three different conditions (low salinity/low temperature, high salinity, or high temperature). The site2 monomers were not constrained and remained structurally flexible during the dissociation process. To ensure convergence in the PMFs, we employed long windows (54 ns) in simulations totalling 52 μs (details provided in the SI; see **Figure Supplement S3** for resulting histograms and PMFs along the dissociation coordinate). According to the free energy difference between the dimerization and dissociation states (ΔG=G_dimer_-G_dissociation_), we estimated the energetic impact from increased salinity and temperature as ΔΔG=ΔG_high salinity or temperature_ -ΔG_293k, 0.15M NaCl_ (**Fig. 3E)**. Notably, high salinity (0.50 M NaCl) or temperature (313 K) decreased the stability of the H-NS_ST_ site2 dimer by 2.2 kcal/mol. H-NS_BA_ displayed a similar sensitivity to temperature but a lower sensitivity to salinity, which destabilized the dimer by 1.5 kcal/mol. Interestingly, our data indicated that H-NS_EA_ was only sensitive to salinity, whereas raising the temperature had little impact on the stability of the H-NS_EA_ site2 dimer. Conversely, H-NS_IL_ only responded to temperature, whereas the increased salinity did not affect the stability of its site2 dimer (ΔΔG ~ 0 kcal/mol). Collectively, our conventional MD simulations and PMF calculations suggested how, on the atomic level, changes in the site2 sequence may alter the sensitivity of the H-NS orthologs to different environmental changes.

### The autoinhibited H-NS conformation is maintained through dynamic electrostatic interactions

In a previous study (*17*), we had shown that melting and dissociation of site2 dimers allow H-NS_ST_ to adapt a closed conformation in which the linker-DNAbd fragment interacts with a negatively charged region on site1 α3 (**Figure Supplement S4A**), and that this auto-inhibitory interaction is incompatible with DNA interactions. However, due to extensive signal broadening of mainly linker amides exchanging with water, our conventional proton-detected NMR analysis based on exchangeable amide H/N-observed correlations did not allow confident mapping of the binding site on the C-terminal region (*17*)(**Figure Supplement S4B**). Herein, we overcame this limitation by using proton-less _13_C-detected NMR analysis to complete the resonance assignment of the linker-DNAbd fragment (**Fig. 4 and Figure Supplement S4C**). These complete carbon chemical shifts allowed us to elucidate the structural mechanism of H-NS_ST_ autoinhibition fully, and, in a second step, to use this understanding to investigate the existence of this closed conformation in the orthologs.

**Figure 4.**
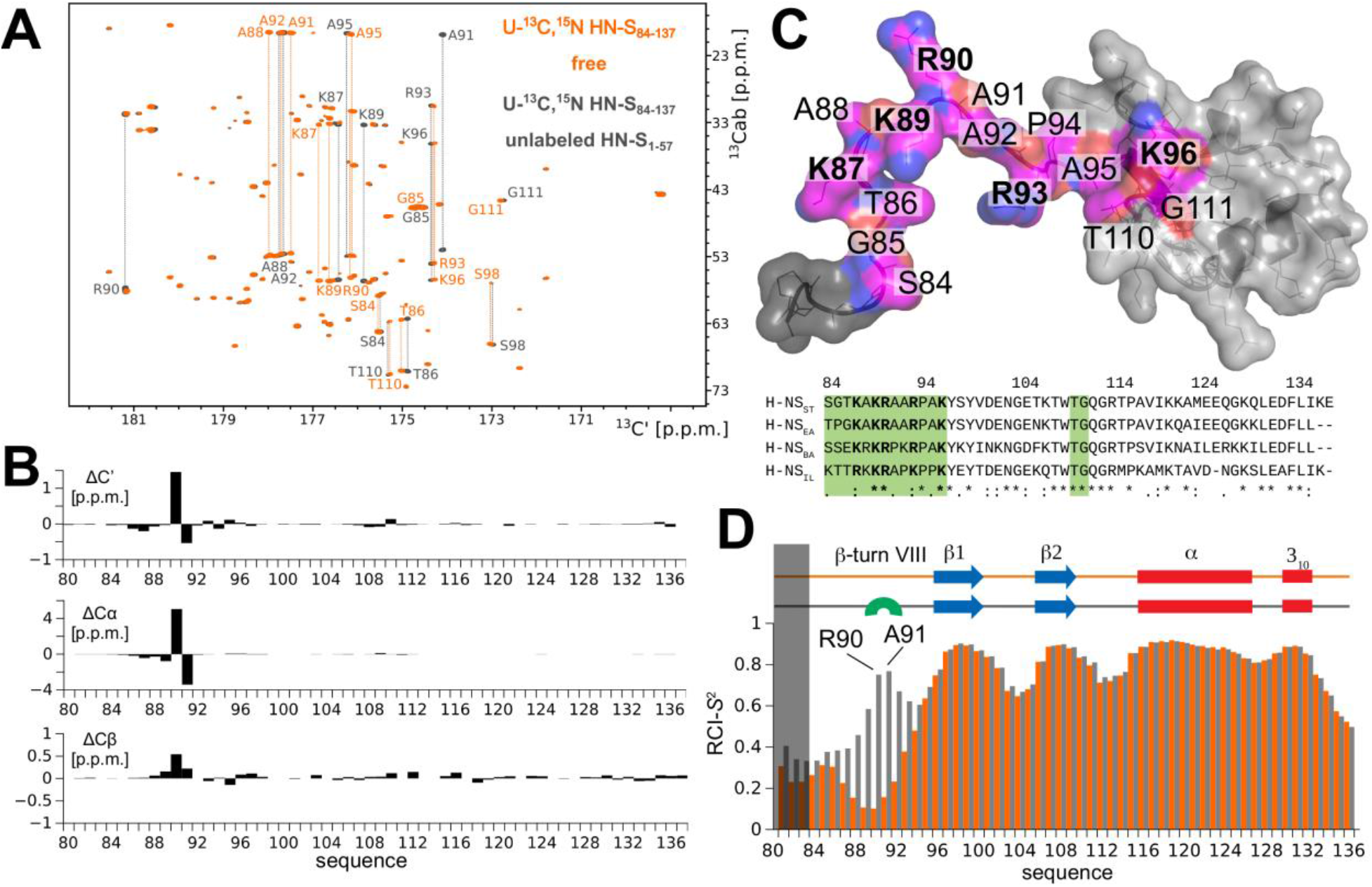
The atomistic details of H-NS auto-inhibition revealed by high-resolution proton-less low-γ detected NMR. **(A)** The 2D CBCACO correlation _13_C-detected spectra of _13_C,_15_N H-NS_ST_Ct (orange) and _13_C, _15_N H-NS_ST_Ct saturated 1:10 (molar) with unlabelled H-NS_1-57_ (dark grey). The OX axis holds all of the _13_C, _15_N H-NS_ST_Ct backbone C’ carbonyl chemical shifts correlated with OY (marked _13_Cab) where each amino acid stripe crosses with its own Cα and Cβ. NMR chemical shift assignments: BMRB IDs 50239 (apo) and 50240 H-NS_ST_ site1–bound. **(B)** *Top panel:* The _13_C chemical shift differences as a function of residue number of H-NS_1-57_ saturated _13_C,_15_N H-NS_ST_Ct and *apo* form. The most marked changes occur in residues K89, R90, A91, and A92 that form a β-turn type VIII. All residues experiencing significant _13_C chemical shift changes upon binding to H-NS1-57 are marked in magenta on the structure of the H-NS_ST_Ct (based on PDBID: 2L93, but extended to contain the full sequence of our construct); positive residues (R+K) are labelled in bold. *Bottom panel:* The sequence alignment of the four selected HN-S orthologs highlighting conserved positively charged linker residues (bold) and residues implicated in binding to site1 (green). **(D)** Secondary structure motifs (red: helix; blue: β-sheet; green: β-turn, present only in the complex) and the RCI-*S*_2_ order parameter (describing the backbone dynamics) of ligand-free (orange) and saturated (gray) H-NS_ST_Ct are shown. For additional details, see **Figure Supplement S4**.

We first titrated unlabelled H-NS_ST_ site1 (residues 1-57) onto the _13_C,_15_N-labelled H-NS_ST_ C-terminal region (Ct_ST_, residues 84-137), comprising the linker (residues 84-93) and DNAbd (residues 94-137) (**Fig. 4A and Figure Supplement S4A**). Motif identification from chemical shifts (*28*) revealed that the interaction promoted the formation of a short type VIII β-turn in residues 89-92 (**Fig. 4B-C**). This sharp turn brings the positively charged linker sidechains K87, K89, R90, R93 closer to each other than in the free state, presumably as a result of pairing them with opposite charges on site1 (*17*) (**Fig. 4B-C**).

To further probe the local dynamics of the polypeptide chain, we determined the random coil index order parameter RCI-*S*_2_ based on the fully assigned _13_C-resonances for each residue for the ligand-free and site1-saturated Ct_ST_. The dynamics of the well-ordered DNAbd domain remained unchanged with or without site1 present, in agreement with its only minor involvement in the auto-association (**Fig. 4D**). Conversely, the linker residues 84-95 were disordered without regular secondary motifs in the absence of site1 (RCI-*S*_2_ < 0.35). Upon addition of site1, the local dynamics decreased, particularly within the stretch of four amino acids K98-R90-A91-A92 (RCI-*S*_2_ > 0.6) that form the type VIII β-turn according to MICS. Nonetheless, the overall RCI-*S*_2_ of the linker remained low, demonstrating that the association with site1 did not substantially restrict the linker’s movements.

Collectively, our analysis established that the autoinhibitory site1:Ct_ST_ association was driven by oppositely charged residues located on site1 and the linker, and involved only a small region of the DNAbd. The resulting intramolecular interaction was maintained through ‘fuzzy’ charge-pairing that did not fix the partners into a structurally stable complex.

### Autoinhibition varies among H-NS orthologs

Having established the detailed autoinhibitory interactions between site1 and the Ct region in H-NS_ST_, we next examined the HNS orthologs. Based on our structural models (initial homology models and models from conventional MD), the electrostatic surface of the Ct was well conserved across all H-NS orthologs (**Fig. 5A**). This level of conservation was expected, given that this region is also required for DNA association (*16*) — a role that needs to be conserved in all H-NS. Conversely, the site1 surface that binds to Ct was not conserved across all orthologs. While H-NS_EA_ was similar to H-NS_ST_ in the overall charge distribution, α3 of H-NS_BA_ showed a distinctly basic surface. H-NS_IL_ displayed an intermediate electrostatic character, with features closer to H-NS_ST_/H-NS_EA_ (**Fig. 5A**). These findings indicated that the stability of the closed conformation varies across orthologs. To test this prediction, we carried out *in vitro* binding experiments using microscale thermophoresis (MST).

**Figure 5.**
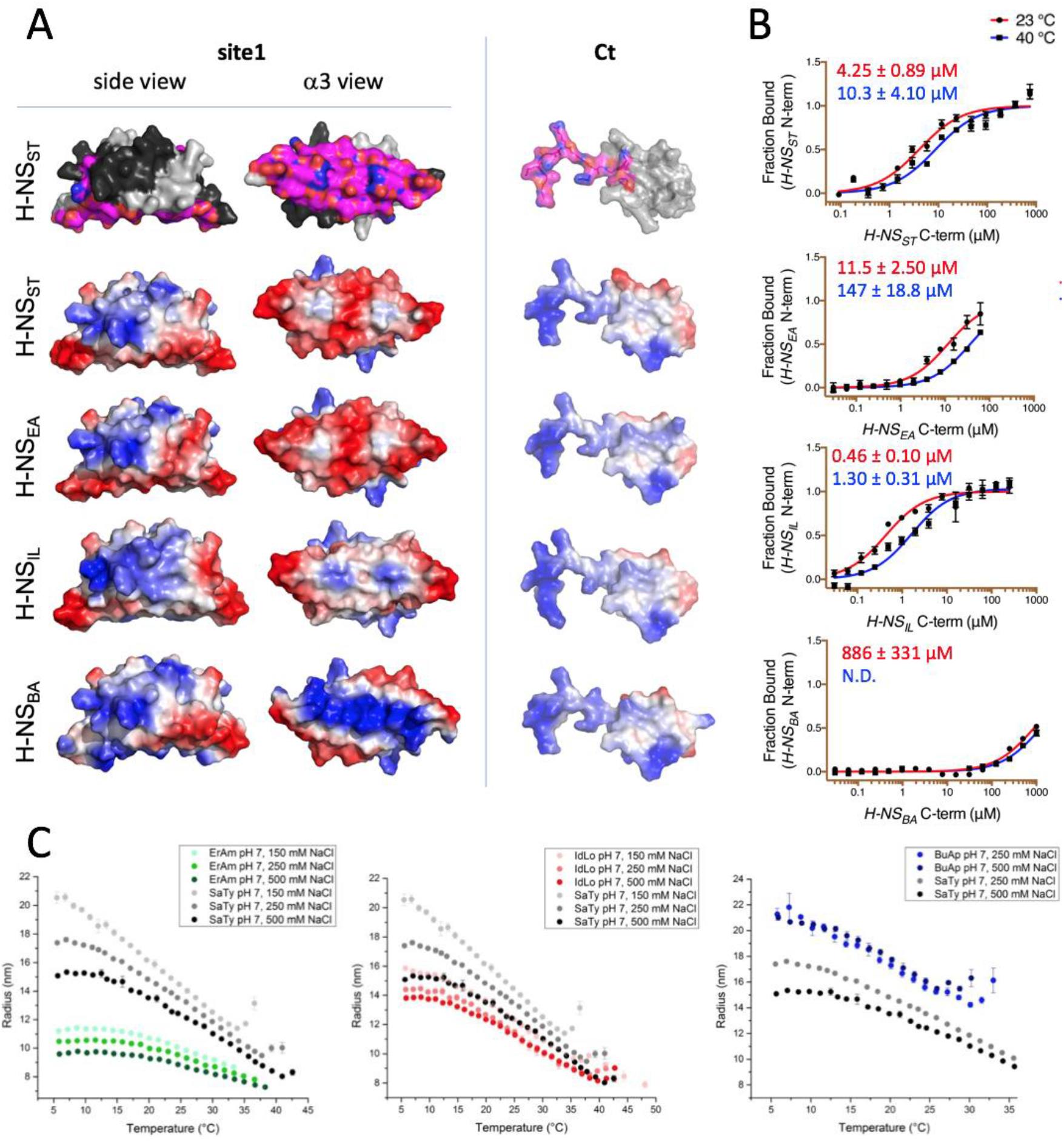
Modeling and *in vitro* analysis of autoinhibition and multimerization of H-NS orthologs. **(A)** *Left and middle panel:* Surface representation of site1 of H-NS orthologs (modeled on *E. coli* H-NS, PDB accession 1NI8) shown as side and α3 (bottom) view. *Right panel:* Ct, comprising the linker-DNAbd fragment, residues 83-137. The top row shows the residues mapped by NMR involved in forming the auto-inhibitory closed conformation ((*17*) and this study). Other rows show the electrostatic surfaces, colorramped from blue (positive) to red (negative) (calculated and visualized by Pymol). **(B)** Microscale Thermophoresis (MST) titrations of unlabelled Ct onto 50 nM of alexa488–labelled site1 at 23 °C (red) and 40 °C (blue). The dissociation constant *K_d_* is color-coded in red (23 °C) and blue (40°C). N.D.: not determined. **(C)** DLS experiments showing changes in hydrodynamic radius (as a proxy of apparent site2 stability) upon changes in salinity and temperature. Data in (B,D) are means ± S.D., *n* = 3.

*In vitro* binding experiments between site1 and Ct corroborated that the strength of the auto-association was similar for H-NS_ST_ and H-NS_EA_ (**Fig. 5B**). H-NS_BA_ has mostly lost its capacity for auto-association, in agreement with its altered site1 surface characteristic. Moreover, H-NS_BA_Ct has a proline residue (P91) in position 3 of the β-turn region, which is highly unfavorable for this secondary structure (*29*). Conversely, the auto-association was tenfold stronger in H-NS_IL_ than in H-NS_ST_, despite a less acidic site1 and despite the presence of a mildly unfavourable proline in β-turn position 4 (P92). Hence, autoinhibition in H-NS_IL_ might include additional and/or different interactions. Increasing the temperature decreased the self-association strength 2–3 fold in H-NS_ST_ and H-NS_IL_, and more than tenfold in H-NS_EA_ (**Fig. 5B**). It also decreased the *K_d_* for H-NS_BA_ to values beyond the measurement range. We concluded that the strength of the autoinhibitory conformation is mostly modulated by the electrostatic surface characteristics of site1, with additional influence from β-turn–breaking proline residues in the Ct, which otherwise preserves its basic characteristic needed for DNA interactions.

### H-NS orthologs show adaptative features *in vitro*

We next experimentally assessed the response of the H-NS orthologs to physicochemical changes using dynamic light scattering (DLS). DLS provides the average hydration radius *R_H_* of the particles in solution, and hence gives a proxy for the tendency of H-NS molecules to form site2-mediated multimers or (still site1-linked) dimers. Thus, the *R_H_* is a convoluted signal of both effects, i.e., the relative strength of site1 multimerization and of the auto-inhibitory conformation (if it exists). We measured the *R_H_* under different salt concentrations and temperatures. As reported previously, H-NS_ST_ showed a clear drop in *R_H_* from 10 to 40 °C (**Fig. 5C**) (*17*). The marked decrease of *R_H_* for curves at 0.15, 0.25, and 0.50 M NaCl indicated a strong inverse correlation between salinity and site2 stability.

All three H-NS orthologs displayed a similar behaviour overall, further supporting that the general mechanism of site2-mediated multimerization and environment sensing was preserved. However, we noted important differences in the orthologs’ response characteristics (**Fig. 5C**): (i) Of the four orthologs, H-NS_ST_ responded most strongly to salinity and temperature, consistent with the broken R54-E74’ salt bridge and large site2 RMSF in our high-salinity or high-temperature simulations. (ii) H-NS_EA_ was less temperature-sensitive and showed weaker multimerization than the other orthologs. Indeed, our simulations suggested that H-NS_EA_ site2 can rearrange the interdimer salt bridge and form either R54-E74’ or R54-D71’ to maintain site2 stability at higher temperatures. (iii) H-NS_BA_ had the highest tendency to multimerize among all the orthologs tested, which might partly be explained by an absence of the autoinhibitory conformation. Compared to H-NS_ST_, our PMF calculations showed a slightly higher sensitivity to temperature and a slightly reduced sensitivity to salinity. Although these tendencies were apparent in our DLS data, these data were also affected by the fact that H-NS_BA_ required more than 150 mM NaCl to stay in solution, but H-NS_BA_ nonetheless aggregated at 30 °C. (iv) H-NS_IL_ showed a decreased sensitivity to salinity compared to H-NS_ST_, as suggested by our computational analysis (i.e., the lack of the site2 K57-R68’ salt bridge, the lack of salt-promoted free energy changes, and the attenuated electrostatic site1 surface). Collectively, our experimental observations revealed significant differences in response to physicochemical parameters, which were in agreement with our predictions based on the molecular features of the H-NS orthologs.

### Conclusions

Environment-sensing through the pleiotropic gene regulator H-NS helps *S. typhimurium* to adapt when it is present inside its host mammal. In a previous study, we had shown that an increase in temperature, and to some extent salinity, dissociates the second dimerization element (site2), which produces two effects: firstly, it impedes synergistic DNA binding of H-NS multimers, and secondly, it allows H-NS to adopt an autoinhibitory conformation where DNA binding residues on the C-terminal linker-DNAbd fragment associate with the N-terminal site1 dimerization domain (*17*). In this study, we confirmed key aspects of this model, namely that site2 is the element that senses changes in physicochemical parameters. We also uncovered additional aspects of this process. In particular, proton-less NMR fully revealed the position and dynamics of the linker-DNAbd residues involved in the autoinhibitory association with site1. The β-turn linker residues 89–91 critical for autoinhibition cannot reach site1 without site2 dissociation (**Figure Supplement S1B**), confirming that the closed autoinhibited conformation is mutually exclusive with H-NS multimerization along DNA. Our NMR analysis also demonstrated that this autoinhibition is achieved at a low entropic cost, maintaining a high flexibility with respect to the exact distribution of the interacting charges on both site1 and the linker-DNAbd fragment. On the one hand, avoiding the entropic penalty helps the autoinhibitory interaction to prevail against the competing DNA association. (Of note, the covalent link between site1 and the Ct will enhance their local concentration and hence their apparent affinity compared to our measurement based on separate domains in **Fig. 5B)**. On the other hand, the fuzziness of the charge-charge interactions facilitates preserving the autoinhibition during bacterial evolution and adaptation.

Based on our refined understanding of the molecular details on the stability and dynamics of site2 and the site1:Ct association of *S. typhimurium* H-NS, we then investigated environment-sensing of H-NS orthologs from bacteria that infect plants, bacteria that are endosymbionts of insects, and bacteria that are presumably free-living in or close to a hydrothermal vent. Across all four orthologs, we observed a conceptually similar response to temperature and salt, both overall and on an atomic level, where salt bridges play key roles. This similarity suggests that environment-sensing in H-NS evolved by adaptation of an ancestral feature, namely the relative instability of the simple site2 helix-turn-helix dimerization motif. However, marked idiosyncrasies in the response of H-NS orthologs suggest that this ancestral feature was adapted to fit the current habitat and lifestyle. Our combined copmutational and experimental structural analysis allowed us to relate the observed *in vitro* features to events on a residual level: in particular, the salt bridge disposition and stability of site2, and the strength of the autoinhibition governed mostly by the electrostatics of site1 helix α3.

Although other factors inside bacteria can modify the *in vitro* behaviour of the isolated protein, it is interesting to consider these idiosyncrasies with respect to the bacteria’s habitats (**Fig. 6**):

i. H-NS_ST_ had the highest sensitivity to temperature and salt, in agreement with the critical role of H-NS_ST_ in helping *Salmonella* adapt its gene expression profile depending on if it is inside or outside a warm-blooded mammal.
ii. In comparison, we found that the response to temperature was markedly attenuated in H-NS_EA_. *E. amylovora* is the causing agent of fire blight, a contagious disease that mostly affects apples and pears (*30*). The reduced sensitivity of H-NS to temperature may reflect the minor importance of this factor in an environment of ambient temperature in temperate climate zones.
iii. *B. aphidicola* is an intracellular symbiont of aphids that is maternally transmitted to the next generation *via* the ovaries (*31*). *B. aphidicola* co-evolved with aphids for more than 150 million years, and despite having the highest sequence identity (61%) to H-NS_ST_ of all orthologs, H-NS_BA_ showed the least conserved features among the orthologs tested, indicating that adaptive evolution was achieved by only minor changes. H-NS_BA_ site2 interactions were stronger than those of other orthologs, and the features promoting the autoinhibitory form were compromised. Hence, H-NS_BA_ may provide a stronger and more robust repression of the genes that it controls. *In vitro*, H-NS_BA_ was the least stable ortholog tested, and had already started to aggregate above 30 °C, in agreement with the fact that *B. aphidicola* cannot survive temperatures of 35°C for extended periods.
iv. Despite having a sequence identity least similar to H-NS_ST_ (41%), H-NS_IL_ maintained an overall similar response profile. However, with an aggregation temperature of 45-50 °C, H-NS_IL_ was the most heat-stable, especially at low pH and high salinity, as expected for a thermophilic and halophilic bacterium. Moreover, autoinhibition was tenfold stronger than in *Salmonella* H-NS and relatively little affected by heat. The natural environment of *I. loiheinsis* (hot hydrothermal fluids venting into cold seawater) provides a temperature range from 4 to 163 °C (*32*), suggesting that temperature-sensing by H-NS_IL_ is biologically relevant. The attenuated response of H-NS_IL_ to salinity might reflect the capacity of *I. loiheinsis* to grow in 20 % (w/v) NaCl medium.

**Figure 6.**
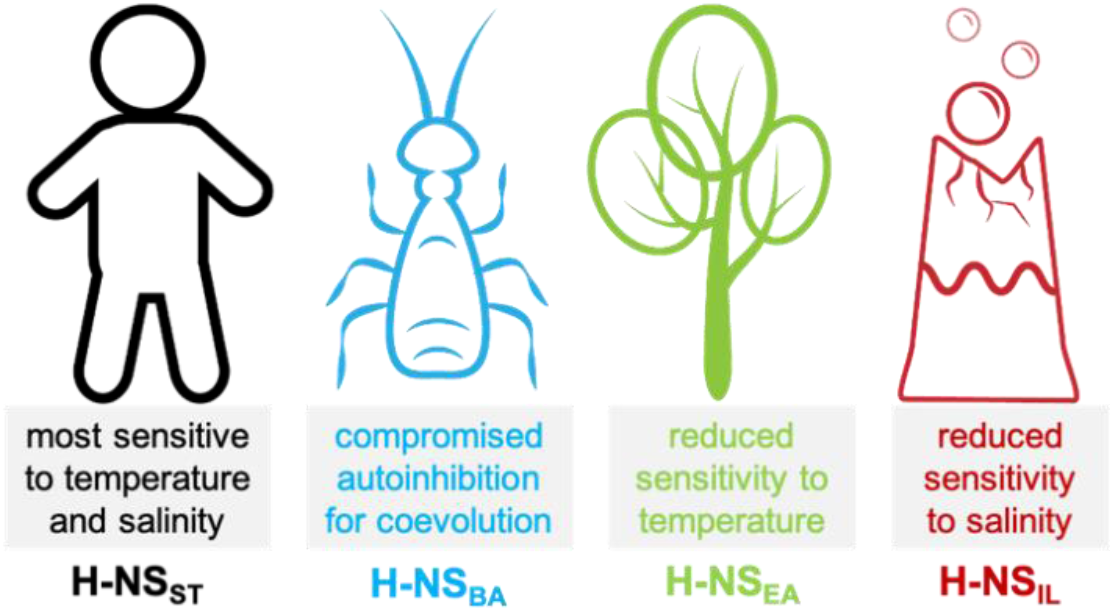
Summary of the most notable adaptations in environment sensing observed for the H-NS orthologs.

Our integrative approach provided atomistic insights on how residue-level substitutions on a protein support adaptation of organisms to different lifestyles. These molecular insights into bacterial adaptative evolution may inspire new strategies to treat related human diseases and to combat antibiotic resistance in the future.

## METHODS

### 1. Simulation Details

Using the CHARMM36 all-atom force field, we performed conventional MD simulations of H-NS tetramers and umbrella sampling of site2 dimers in GROMACS. For DLS and MST, recombinant protein production and measurements were adapted from (*17*). However, we fluorescently labelled site1 for DLS, instead of the Ct. For NMR, _13_C,_15_N-labelled *S. typhimurium* H-NS84-137 was expressed in minimal M9 media with 5 g/L of U-_13_C glucose and 1 g/L of _15_NH_4_Cl salt. Proton and low-γ detected high-resolution NMR spectroscopy was carried out on a 700-MH Bruker NEO spectrometer equipped with a 5-mm cryogenic TXO direct observe _15_N,_13_C-optimized probe at 25 oC. Further details are shown as below.

#### 1.1 Model preparation

We built our homology model of full-length H-NS (UniprotID: P0A1S2) based on orthologues in SwissModel (*33*) with the templates for the dimerization domain (PDBID: 3NR7) and the DNA-binding domain (PDBID: 2L93). The site2 dimer models were initiated in an anti-parallel configuration, while the tetramer models were constructed according to the crystal packing (PDBID: 3NR7). Maestro (Schrödinger, Inc.) was used to construct the full-length model from different domains.

#### 1.2 Simulation setup

Our simulations were carried out by GROMACS (*34*) (MD simulations of tetramers and Potential of Mean Force (PMF) simulations of site2 dimers). All the models were solvated in a TIP3P water box, with counterions to neutralize the charges and additional NaCl for the desired salinity. Each tetramer system contains ca. 33,000 TIP3P water molecules, counter ions, and 150 or 500 mM NaCl, totaling ca. 100,000 atoms in a periodic box 13 ×9×9 nm3. All simulations were performed following a minimization, 250 ps equilibration in the NVT and NPT ensemble with Berendsen temperature and pressure coupling, and a production stage NPT (293 or 313 K, 1 bar). The CHARMM36 force field (*35*) was used with the cmap correction. The particle mesh Ewald (PME) technique (*36*) was used for the electrostatic calculations. The van der Waals and short-range electrostatics were cutoff at 12.0 Å with switch at 10 Å.

The PMF simulations were carried out with the MD program GROMACS (*34*) using the umbrellaing sampling (US) technique. The CHARMM36 force field was also used. Each site2 monomer of the center of mass (COM) distance was chosen as the disassociation pathway and used for enhanced sampling. After 500 ps equilibrium with the NPT ensemble, initial structures for windows along the reaction coordinates were generated with steered MD. In the steered MD simulation, one chain was pulled away along in the direction of increasing the COM distance with force constant of 12 kcal mol_-1_ Å_-2_, until the COM distance reached 25 Å. The windows were taken within a range of 0-25 Å. The umbrella windows were optimized at the 0.3 Å interval to ensure sufficient overlap. There are about 80 windows per simulation, and each window was simulated with force constant of 1.2 kcal mol_-1_ Å_-2_. All PMF simulations converge in 54 ns per window (**Supplementary Fig. S1**).

#### 1.3 Computational Data analysis

All the data analyses were carried out in GROMACS, VMD Tcl scripts, and in-house Python programs. In particular, root-mean-square fluctuations (RMSF), root-mean-square deviation (RMSD), salt-bridge interaction, polar interaction, and hydrophobic interaction were analyzed in VMD (*37*) For salt-bridges, the O and N atoms in the charged residues (Arg, His, Lys, Asp, and Glu) were used with a distance cut-off of 4.5 Å. For the polar interaction, the atoms in the side chains with the partial charge cutoff (> 0.3 unit for a polar contact) were used with a distance cutoff of 4.5 Å. The cut-off for classifying hydrophobic interactions was 6.0 Å between the C atoms of the hydrophobic residues. The relative percentage (*P*) of hydrophobic contacts is defined as **Eq. 1**.

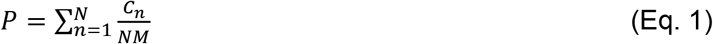

N is the total number of frames, *C_n_* is the number of hydrophobic contacts at frame *n*, and *M* is the largest value in the series of *C_n_*. The PMFs were determined using the Weighted Histogram Analysis Method (WHAM) (*38*) implemented in GROMACS. All the visualization was performed with python, VMD, Pymol (Schrödinger, Inc.), and Maestro (Schrödinger, Inc.).

### 2. Experimental Details

#### 2.1 Protein production

S. typhimurium H-NS_1-57,C21S_ and H-NS_84-137_ were cloned and produced as described previously (*17*). *E. amylovora* (H-NS_1-57_, H-NS_82-134_), *B. aphidicola* (H-NS_1-57_, H-NS_84-135_) and *I. loihiensis* (H-NS_1-57_, H-NS_85-138_) genes were individually cloned into pGEX6P-1, expressed and purified as described previously (*17*). For the high-resolution nuclear magnetic resonance (NMR) studies the uniformly double _13_C,_15_N-labelled *S. typhimurium* H-NS_84-137_ (with additional GPLG residues before S84) was expressed in minimal M9 media with 5g/L of U-_13_C glucose and 1g/L of _15_NH_4_Cl salt. The unlabeled *S. typhimurium* H-NS_1-57_ *N*-terminal domain was expressed and purified as before (*17*). The final NMR buffer was 50 mM NaCl, 2% (*v/v*) D_2_O, 20 mM Bis-TRIS at pH 6.5 and 0.002% NaN_3_.

#### 2.2 Dynamic Light Scattering

For DLS measurements, H-NS from *Salmonella typhimurium, Erwinia amylovora, Buchnera aphidicola* and *Idiomarina loihiensis* were expressed as N-terminal mCherry fusion proteins with an N-term His tag in *E. coli* BL21 using the expression vector pET28b. The linker sequence SAGGSASGASG was inserted between mCherry and H-NS proteins to avoid steric clashes in the dimer. Bacteria were grown in LB medium, induced with 1mM IPTG at 25 °C overnight. Cells were harvested and resuspended in lysis buffer (50 mM Tris pH8, 500 mM NaCl, 10 mM Imidazole with addition of lysozyme, DNase I and 1% triton X-100) and lysed by mild sonication. Proteins and bacterial membranes were separated by centrifugation (30 min, at 15,000 *×g*) and the supernatant was applied to Ni-NTA beads (Qiagen) for 2h. The column was washed thoroughly with 50 mM Tris pH8, 500 mM NaCl, 10 mM Imidazole and protein was then eluted with 50 mM Tris pH8, 500 mM NaCl, 400 mM Imidazole, 1 mM DTT. After dialysis in 50 mM HEPES pH7.4, 300 mM NaCl, 0.5 mM TCEP, eluted protein was further purified by ion-exchange chromatography using either MonoQ or MonoS column (GE) in the same buffer. Protein multimerization was observed in combination of different salt (150, 250 and 500 M NaCl) and pH (6, 7 and 8) conditions. For this, 100 mM MES, MOPS and HEPES buffers were used, with proteins at concentrations ranging from 125 to 500 μM, in a final volume of 100 μL. Dynamic light scattering measurements were performed in 96-well plates (Greiner) using a DynaPro plate reader-II (Wyatt Technologies). A triplicate of three wells was measured for every sample with 5 acquisitions of 5 s for every well. The machine was cooled with gaseous nitrogen, with a starting temperature of 5 T, followed by an increase to 60 °C at a ramp rate set so that each well is measured every 1°C. Data were analyzed with DYNAMICS software (Wyatt Technologies) as Temperature Dependence and exported for further fitting on Origin software using a Logistic Fit. The presented results are mean values with standard error mean determined from the triplicate sample.

#### 2.3 Proton and low-γ detected high-resolution NMR spectroscopy

All NMR measurements were done on 700 MH Bruker NEO spectrometer equipped with 5 mm cryogenic TXO direct observe _15_N,_13_C-optimized probe at 25°C. The sequence specific backbone resonance assignments of visible H/N correlations on 1H-detected spectra of *S. typhimurium* H-NS_84-137_ protein at 200 μM concentration in *Apo* and H-NS_1-57_ saturated (1.5 mM) forms were achieved with classical set of triple resonance experiments, *i.e*. HNCA, HncoCA, HNCO, HNcaCO, HNCACB, CBCAcoNH (*39*) and previously published assignments (*40*). The 100% complete sets of Cα, Cβ and C’ resonances for *Apo* and H-NS1-57 saturated (1.5 mM) forms covering the entire protein sequence, together with the residues not visible on H/N correlation 1H-detected experiments (due to amide exchange with water) were achieved with intra-residue 2D (H)CACO (*c_hcaco_ia3d*, 16 scans) and (H)CACBCO (*c_hcbcaco_ia3d*, 32 scans) supported with sequential (H)CANCO (*c_hcanco_ia3d*, 96 scans) _13_C-detected experiments (*41*). The low-γ, so _13_C-detected experiments mentioned above were started with 1H-excitation in order to enhance the sensitivity and recorded in in-phase and anti-phase (IPAP) mode for the virtual decoupling. All spectra were processed in NMRpipe and analyzed in CARA and Sparky software. The random-coil-index order parameters RCI-*S*_2_ and secondary motifs, like β-turn, for *Apo* and H-NS_1-57_ saturated (1.5 mM) forms were determined from complete lists of C_α_, C_β_ (except glycines), N, C’ chemical shifts with the TalosN and MICS programs, respectively. NMR chemical shift assignments for the H-NS_ST_Ct in its apo and H-NS_ST_ site1–bound states are deposited at the BMRB with the IDs 50239 and 50240, respectively.

#### 2.4 Micro Scale Thermophoresis (MST) for protein-protein interactions

H-NS (residue 1-57) from *S. typhimurium, E. amylovora, B. aphidicola* and *I. loihiensis* were individually labeled N-terminally with fluorescent Alexa488-TFP (Thermo Scientific) and then unlabeled C-term of those proteins were titrated against Alexa488 labeled N-term correspondingly and the final results were plotted as described previously (*17*).

## ACKNOWLEDGEMENTS

Research by UH, VK, FH, AK, ML, LJ, and SA reported in this work was supported by the King Abdullah University of Science and Technology (KAUST) through the baseline fund and the Award No FCC/1/1976-25 from the Office of Sponsored Research (OSR); XZ was partially supported by the ACS Petroleum Research Fund (58219-DNI); CL, JMR, and JL were partially supported by the National Institutes of Health award (R01GM129431). We acknowledge support from the KAUST Bioscience and Imaging core laboratories and the computational resources from the Vermont Advanced Compute Core (VACC) and the Anton supercomputer in Pittsburgh Supercomputing Center (PSC), and thank M. Cusack (KAUST Research Support Services) for editorial help.

## FIGURE SUPPLEMENTS

**Figure 2 – Figure Supplement S1.** The umbrella histogram and convergency to simulate the HNS orthologs site2 dimer at different simulation conditions: 293 K, 0.15 M NaCl (Green), 293 K, 0.50 M NaCl (Blue), and 313 K, 0.15 M NaCl (Red). Each PMF subplot contains a series of 5 curves from 38 to 54 ns/window (4 ns increment), demonstrating convergence.

**Figure 3 – Figure Supplement S2.** Representative H-NS monomers (with site2 alignment) from the final snapshots (at 200 ns) of the tetramer simulations under different conditions: 293 K, 0.15 M NaCl (green), 293 K, 0.50 M NaCl (cyan), and 313 K, 0.15 M NaCl (red). To highlight the linker region in α3, we use a pale color for the rest of the backbone in each protein. This figure shows that while H-NS proteins are mostly “straight” at 293 K, 0.15 M NaCl (green), bending of α3 is common through all H-NS orthologs under the high-temperature or high-salinity condition. The most significant bending is found in H-NS_ST_.

**Figure 3 – Figure Supplement S3.** The umbrella histogram and convergency to simulate the HNS orthologs site2 dimer at different simulation conditions: 293 K, 0.15 M NaCl (Green), 293 K, 0.50 M NaCl (Blue), and 313 K, 0.15 M NaCl (Red). Each PMF subplot contains a series of 5 curves from 38 to 54 ns/window (4 ns increment), demonstrating convergence.

**Figure 4 – Figure Supplement S4.** The low-γ _13_C-detected experiments unveil the molecular details of highly dynamic and solvent exposed residues elusive for classical _1_H-detected approaches. The panel **A** shows the _1_H-_15_N HSQC spectrum of 200 uM _13_C,_15_N-labelled *S. typhimurium* H-NS_84-137_ protein in *Apo* (orange) and H-NS_1-57_ saturated (1.5 mM) form (dark gray). The residues that are missing from H/N correlation 1H-detected experiments are depicted in panel **B** with black and green mapped on the NMR solution structure of H-NS_91-137_ (PDB id 1HNR; the lowest energy structure). The flexible residues of S84-A91 were added together with GPLG artificial residues left after the tag cleavage. The only tryptophan side chain Hε/Nε imidazole correlation is marked W109 in green. The **C** panel presents the overlay of 2D (H)CACO and (H)CANCO _13_C-detected experiments and sequential walk for two regions, G80-A91 marked black and T110-G111 marked green, that are not detected on **A**. A complete sequential carbon walk can be done for the entire protein H-NS_84-137_ sequence.

**Figure 2 – Figure Supplement S1.**
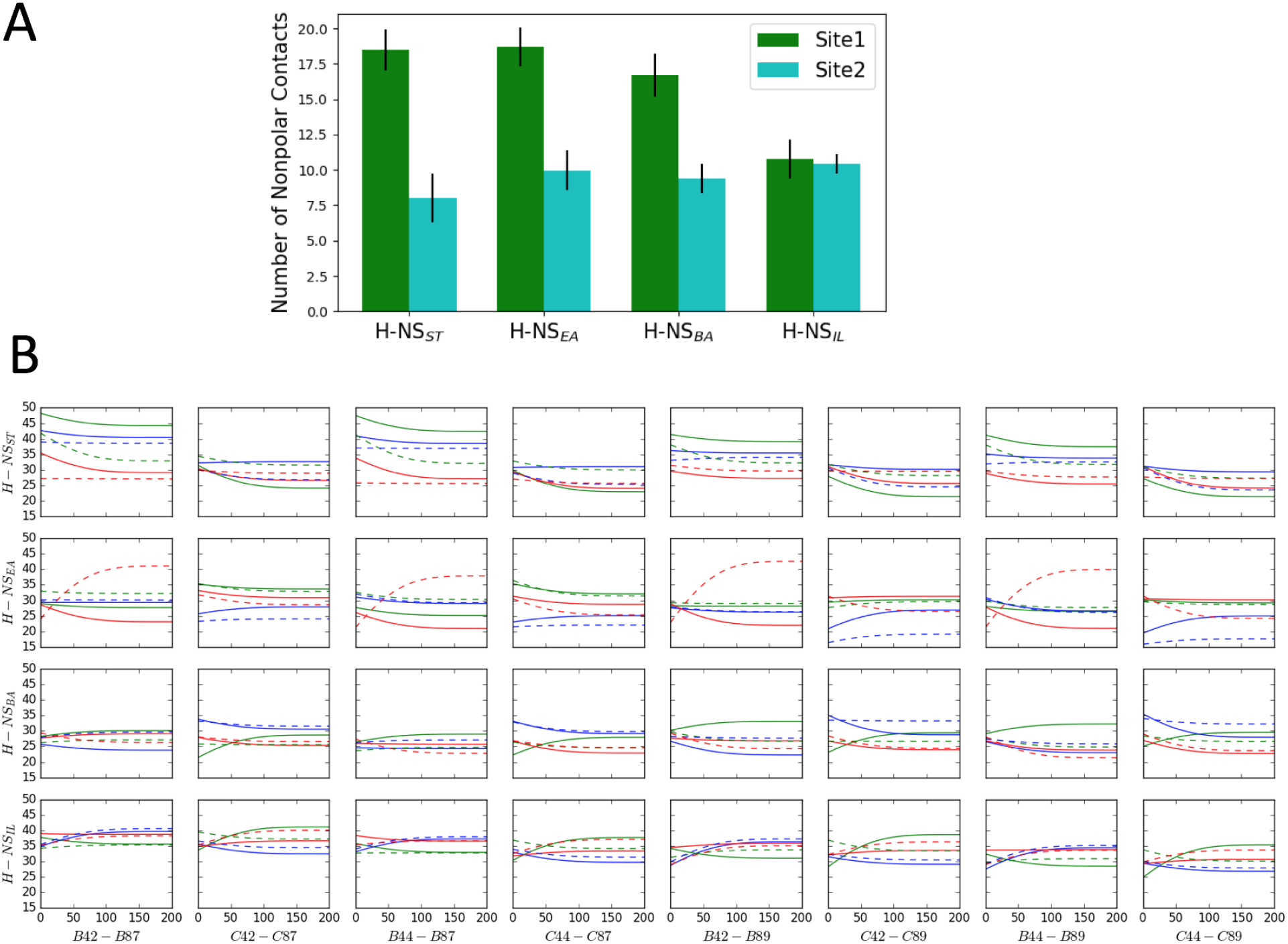
(A) Comparison of nonpolar contacts of H-NS orthologues at 293K and 0.15M NaCl, using the last 100 ns in the MD simulations. (B) The separation distance between residue 42/44 and residue 87/89 (Y axis in Å) versus time (X axis in nanosecond). The separation distance was measured as the C_α_-C_α_ distance in the tetramer model (within chain B/C, which were modelled as full length). The color scheme annotates different simulation conditions: 293 K, 0.15 M NaCl (Green), 293 K, 0.50 M NaCl (Blue), and 313 K, 0.15 M NaCl (Red). For clarity, we show the smoothed data of two replicas for each system (solid and dash lines respectively). Since complete unfolding of site2 was not observed in these simulations (presumably due to the short timescale and the difficulty of sampling), these plots indicate a minimum separation of 15 to 20 Å between the N and C termini, demonstrating that site2 has to unfold to allow closer contacts between the termini.

**Figure 3 – Figure Supplement S2.**
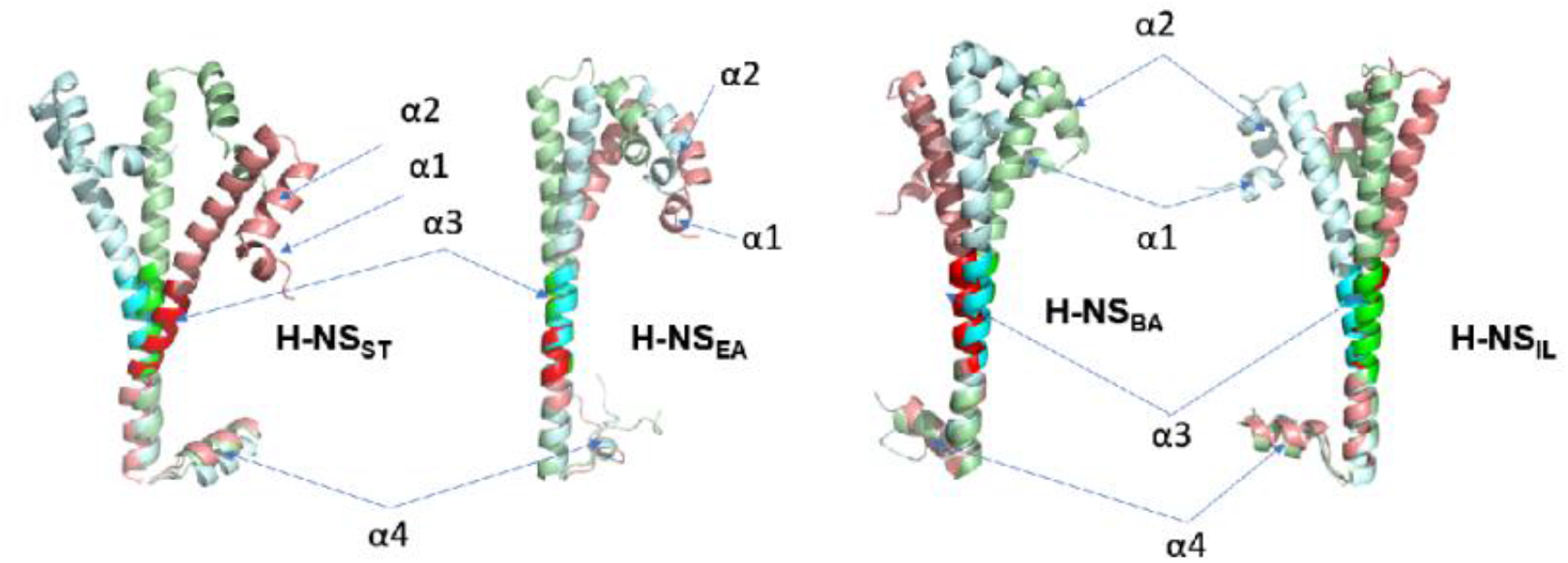
Representative H-NS monomers (with site2 alignment) from the final snapshots (at 200 ns) of the tetramer simulations under different conditions: 293 K, 0.15 M NaCl (green), 293 K, 0.50 M NaCl (cyan), and 313 K, 0.15 M NaCl (red). To highlight the linker region in α3, we use a pale color for the rest of the backbone in each protein. This figure shows that while H-NS proteins are mostly “straight” at 293 K, 0.15 M NaCl (green), bending of α3 is common through all H-NS orthologs under the high-temperature or high-salinity condition. The most significant bending is found in H-NS_ST_.

**Figure 3 – Figure Supplement S3.**
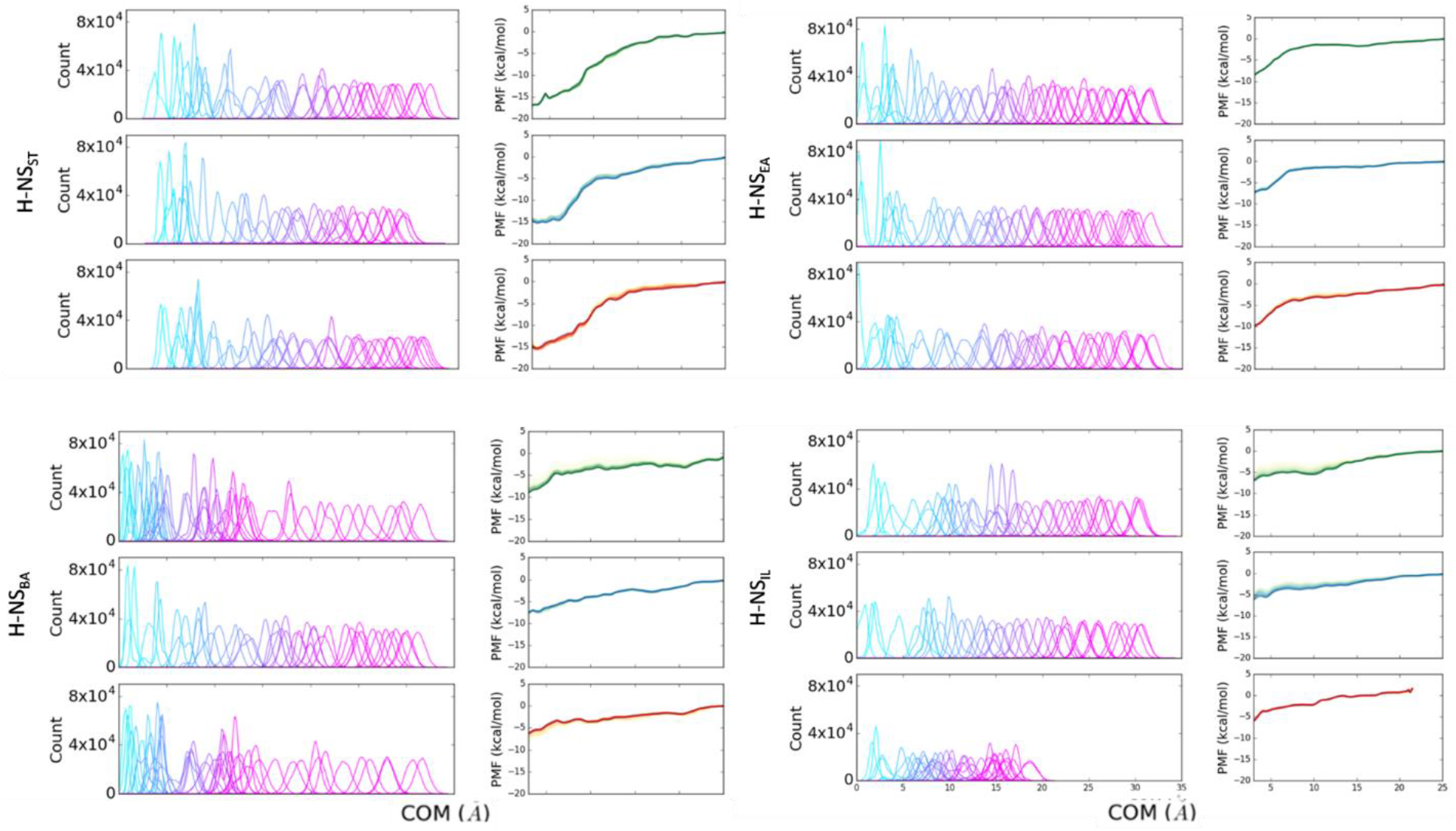
The umbrella histogram and convergency to simulate the H-NS orthologs site2 dimer at different simulation conditions: 293 K, 0.15 M NaCl (Green), 293 K, 0.50 M NaCl (Blue), and 313 K, 0.15 M NaCl (Red). Each PMF subplot contains a series of 5 curves from 38 to 54 ns/window (4 ns increment), demonstrating convergence.

**Figure 4 – Figure Supplement S4.**
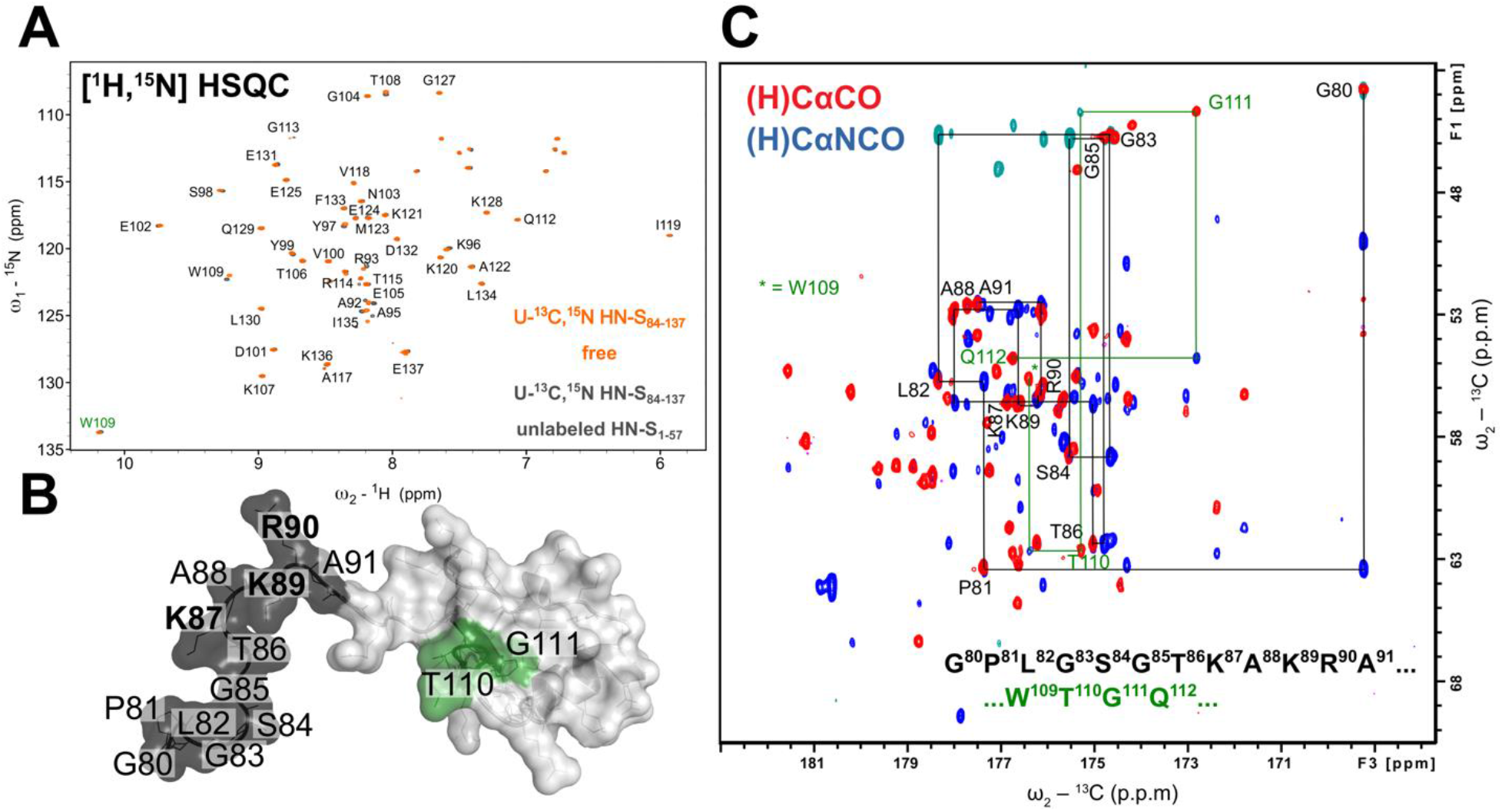
The low-γ ^13^C-detected experiments unveil the molecular details of highly dynamic and solvent exposed residues elusive for classical ^1^H-detected approaches. The panel **A** shows the ^1^H-^15^N HSQC spectrum of 200 uM ^13^C,^15^N-labelled *S. typhimurium* H-NS_84-137_ protein in *Apo* (orange) and H-NS_1-57_ saturated (1.5 mM) form (dark gray). The residues that are missing from H/N correlation ^1^H-detected experiments are depicted in panel **B** with black and green mapped on the NMR solution structure of H-NS_91-137_ (PDB id 1HNR; the lowest energy structure). The flexible residues of S84-A91 were added together with GPLG artificial residues left after the tag cleavage. The only tryptophan side chain Hε/Nε imidazole correlation is marked W109 in green. The **C** panel presents the overlay of 2D (H)CACO and (H)CANCO ^13^C-detected experiments and sequential walk for two regions, G^80^-A^91^ marked black and T^110^-G^111^ marked green, that are not detected on **A**. A complete sequential carbon walk can be done for the entire protein H-NS_84-137_ sequence.

